# Targeted delivery of flagellin by nebulization offers optimized respiratory immunity and defense against pneumococcal pneumonia

**DOI:** 10.1101/2024.06.17.599391

**Authors:** Mara Baldry, Charlotte Costa, Yasmine Zeroual, Delphine Cayet, Jeoffrey Pardessus, Daphnée Soulard, Frédéric Wallet, Delphine Beury, David Hot, Ronan MacLoughlin, Nathalie Heuzé-Vourc’h, Jean-Claude Sirard, Christophe Carnoy

## Abstract

Novel therapeutic strategies are urgently needed to combat pneumonia caused by *Streptococcus pneumoniae* strains resistant to standard-of-care antibiotics. Previous studies have shown that targeted stimulation of lung innate immune defenses through intranasal administration of the Toll-like receptor 5 agonist flagellin, improves the treatment of pneumonia when combined with antibiotics. To promote translation to the clinic application, this study assessed the direct delivery of flagellin to the airways through nebulization using a vibrating mesh nebulizer in mice. Intranasal delivery achieved approximately 40% lung deposition of the administered flagellin dose, whereas nebulization yielded less than 1%. Despite these differences, nebulized flagellin induced a transient activation of lung innate immunity characterized by cytokine/chemokine production and neutrophil infiltration into airways analogous to intranasal administration. Furthermore, inhalation by nebulization resulted in an accelerated resolution of systemic pro-inflammatory responses. Lastly, adjunct therapy combining nebulized flagellin and amoxicillin proved effective against antibiotic-resistant pneumococcal pneumonia in mice. We posit that flagellin aerosol therapy represents a safe and promising approach to address bacterial pneumonia within the context of antimicrobial resistance.

## 1. Introduction

*Streptococcus pneumoniae*, commonly known as pneumococcus, typically inhabits the human nasopharynx but can cause severe and often fatal diseases upon invasion of the host, such as pneumonia [1]. While vaccines prevent pneumococcal pneumonia, they target only a fraction of the nearly 100 serotypes identified thus far [2]. Consequently, there has been a rise in infections caused by serotypes not included in the current available vaccines, many of which exhibit antimicrobial resistance (AMR) [3, 4]. Resistance of *S. pneumoniae* to β-lactam antibiotics such as amoxicillin (the standard of care for pneumococcal pneumonia) as well as macrolides, is a major concern worldwide with the prevalence of resistance likely exceeding 20% in Europe alone [5]. Given the slow pace of new antimicrobial innovation and discovery [6], it is imperative to explore alternative therapeutic strategies to treat pneumococcal pneumonia otherwise untreatable with antibiotics.

Host-immune directed therapy, also referred to as immunotherapy, is a promising alternative strategy to antibiotics aimed at either activating antimicrobial immunity or modulating inflammation [7]. Toll-like receptors (TLRs) are pivotal pattern-recognition receptors involved in orchestrating the antimicrobial immune defenses. TLRs can recognize pathogen-associated or damage-associated molecular patterns and are widely expressed, particularly in immune protein of the bacterial flagellum, demonstrates notable immunotherapeutic potential against respiratory infections [9]. Flagellin is highly soluble and can be readily formulated in a saline solution for delivery in human, including inhalation. TLR5 is expressed by airway epithelial cells that are recognized as the principal drivers of TLR5-mediated protection. These cells orchestrate the production of cytokines, chemokines, and antimicrobial peptides, and facilitate the recruitment and activation of immune cells within the airways, particularly neutrophils [10-When combined with oral amoxicillin or intraperitoneal sulfamethoxazole, inhaled flagellin demonstrates synergistic effects, leading to a substantial decrease in pneumococcal load within the lungs of infected mice, otherwise not achieved with antibiotic or flagellin monotherapy [15, 16]. This combined treatment enables a reduction in the antibiotic dosage required to achieve a comparable therapeutic outcome observed with antibiotic. More importantly, the synergistic effect is also seen when the mice are infected with amoxicillin-resistant pneumococcus [15]. Additionally, adjunct flagellin therapy to antibiotics attenuates the selection of antibiotic resistant strains in a co-infection model emphasizing the relevance of this approach in combating AMR [17]. Another example of TLR agonist application in respiratory For treating infections of the lower respiratory tract, *i.e.,* pneumonia, direct delivery to the lungs is obvious, matching the delivery route and the pathogen location. The method of delivering therapeutics to the lungs depends on the clinical situation, the drug nature and its formulation, and it is crucial to deliver a therapeutic dose of the drug to the target site [22–26] While intranasal administration is a common method for targeting therapeutics into conducting airway in animal studies, it is not adapted to target the lungs, in particular the deep lungs, in humans [27]. Nebulization delivery, which produces an aerosol, stands out as the most efficient approach for administering drugs and biologics as liquid forms, into the lungs in patients under both spontaneous breathing and mechanical ventilation [23–25]. Vibrating mesh nebulizers allow for fast, effective and user-friendly drug delivery to the airways [28–30]. Moreover, they are usually more appropriate for biologics, limiting adverse effects during the aerosol generation [24, 26, 31]. Adapting flagellin to aerosol delivery is an important step towards the development of clinical investigations.

Here, we describe the use of a custom-made nebulization chamber for nose-only delivery of aerosols in mice using a vibrating mesh nebulizer, and studied the fate and activity of nebulized flagellin (hereafter referred to as FLAMOD), the activation of lung innate immunity, and the therapeutic efficacy in combination with antibiotics against pneumococcal pneumonia. Comparing with intranasal administration as a benchmark, our study reveals that nebulizer-mediated delivery of FLAMOD demonstrates comparable or superior efficacy in promoting lung immunity and protection, despite lower amounts being delivered to the lung. Moreover, it induces a less pronounced and enduring impact on systemic pro-inflammatory responses.

## 2. Materials and Methods

### 2.1 Bacterial strains

*Streptococcus pneumoniae* serotype 1 (clinical isolate E1586) was obtained from the National Reference Laboratory—Ministry of Health, Uruguay [12]. *Streptococcus pneumoniae* serotype 19F (clinical isolate LILPNEUHC 19F) was obtained from the University Hospital of Lille (Frédéric Wallet, CHU Lille, France); it is a multidrug-resistant clinical isolate from a bacteremia patient, and is resistant to amoxicillin (minimal inhibitory concentration, MIC_AMX_ = 1.5 μg/ml), erythromycin and tetracycline, and has intermediate resistance to cefotaxime and ceftriaxone. Working stocks were prepared as described previously [12]. Briefly, fresh colonies grown on blood-agar plates were incubated in Todd Hewitt Yeast Broth (Sigma-Aldrich, Saint-Louis, MO) at 37^◦^C until the OD600nm reached 0.7–0.9 units. Cultures were stored at −80^◦^C in broth + glycerol 12% (vol./vol.) for up to 3 months. For infection, working stocks were thawed and washed with sterile Dulbecco’s Phosphate-Buffered Saline (PBS, Gibco, Grand Island, NY) and diluted to the appropriate concentration. The number of bacteria (as colony forming units [CFUs]) was confirmed by plating serial dilutions onto 5% sheep blood agar plates and incubating at 37°C for 18 h with 5% CO_2_.

### 2.2 FLAMOD preparation

Flagellin was produced in *Escherichia coli* by the Vaccine Development department at Statens Serum Institut, Denmark. The custom-designed flagellin used in the present study, *i.e.,* FLAMOD (recombinant flagellin FliC_Δ174-400_ as described in Nempont et al 2008 [32] harboring one extra amino acid at the N terminus) derives from *Salmonella enterica* serovar Typhimurium FliC. FLAMOD was purified by filtration and chromatography and resuspended in NaPi buffer: 10 mM phosphate, 145 mM NaCl, polysorbate 80 0.02 % (w/v) pH 6.5. Immunostimulatory activity was validated using the HEK-Dual™ hTLR5 cells assay (Invivogen). The endotoxin content in the protein preparation was assessed with a *Limulus* assay (Pyrochrome, kinetic LAL assay from Associates of Cape Cod Inc.).

### 2.3 Mouse treatment and infection

All experiments complied with institutional regulations and ethical guidelines, were approved by Institutional Animal Care and Use Committee (E59-350009, Institut Pasteur de Lille; Protocol APAFIS#16966 201805311410769_v3) and were conducted by qualified personnel. Female BALB/cJRj mice, male C57BL/6JRj mice (6–8 weeks old) (Janvier Laboratories, Saint Berthevin, France) and female and male *Tlr5^-/-^* (varying ages, breed in-house on C57BL/6JRj background) were maintained in ventilated cages (Innorack® IVC Mouse 3.5) and handled in a laminar flow biosafety cabinet (Class II Biohazard, Tecniplast).

Intranasal treatment was performed with 2.5 μg or 30 μg FLAMOD in 30 μl NaPi buffer that was administered intranasally under light anesthesia via isoflurane inhalation (Axience, Pantin, France). The dose of 2.5 µg of FLAMOD has been previously demonstrated protective in pneumococcal infections while the dose of 30 µg corresponds to the amount of FLAMOD used for the biodistribution analysis. Control animals (mock treated) received intranasal NaPi buffer alone. For FLAMOD administration using the Aerogen^®^ Solo vibrating mesh nebulizer (Aerogen, Ireland) in combination with ProX controller (Aerogen, Ireland), a custom-made chamber was used (EuroBioConcept, Bonneuil sur Marne, France). One milliliter of NaPi buffer or FLAMOD at 250 µg/mL was administered for 4-5 minutes to vigilant mice using the Aerogen^®^ Solo nebulizer (**Supplementary Figure 1**). For intravenous administration, FLAMOD was injected retro-orbitally at concentrations of 10 or 30 µg per mouse prepared in 200 μl PBS. After treatment, mice were sacrificed via the intraperitoneal injection of 5.47 mg sodium pentobarbital in 100 μl PBS. Broncholalveolar lavage (BAL) fluids were obtained by intratracheal injection of 2 × 1 ml PBS supplemented with protease inhibitors (Roche). Serum was collected by intracardiac puncture and separated from whole blood in Z-Gel microtubes (Sarstedt). BAL and serum samples were stored at −20^◦^C prior to analysis.

Prior to infection, the mice were anesthetized via the intraperitoneal injection of 1.25 mg (50 mg/kg) ketamine plus 0.25 mg (10 mg/kg) xylazine in 250 μl of PBS. For primary infections with the serotype 1 isolate E1586 in BALB/c mice, 2×10^6^ CFU were inoculated intranasally in 30 μl PBS, as described previously [16]. For the pneumococcal superinfection, C57BL/6 mice were used as previously described [16, 33]. In this model, animals were first infected i.n.with 50 plaque-forming units (PFU) of the pathogenic, murine-adapted H3N2 influenza A virus strain Scotland/20/74 in 30 µl of PBS. Seven days later, i.n. *S. pneumoniae* infection was induced with 1×10^5^ CFU of the serotype 19F strain in 30 µl of PBS. Mice were treated 12 h post-infection with intranasal or nebulized FLAMOD and with intragastric administration of 5 μg or 1 mg of amoxicillin (Clamoxyl for injection, GlaxoSmithKline) in 200 μl of water. Mock treated control animals received NaPi buffer and water. After 24 h infection, lungs and spleen were homogenized with an UltraTurrax homogenizer (IKA-Werke, Staufen, Germany), and viable bacterial counts were determined on blood agar plates.

### 2.4 FLAMOD-specific ELISA

Maxisorp 96 well (Nunc 442404) plates were coated with 50 µl of capture anti-FLAMOD monoclonal antibody (Mab, 9H10-R2-2E8 mouse IgG1 - lot vt220802-7034) at 5µg/ml in D-PBS 1X (PBS, Gibco 14190-094) and incubated overnight at 4°C. The plates were washed 3 times with PBS/Tween 20 0.05% (PBS-T) and saturated for 2 h at room temperature (RT) with 300 µl per well ELISA/ELISPOT diluent 1X (Invitrogen 00-4202-56, 5X solution diluted in sterile water Versylene Fresenius B230531). The plates were washed again 3 times with PBS/T. Serum samples were diluted 1:5 in ELISA /ELISPOT diluent (i.e., 20% final ratio of mouse serum), BAL samples 1:200 and 50 μl were distributed per well in the microplates. To generate a calibration curve, FLAMOD (lot 4182.5) was diluted from 100 ng/ml to 0.78 ng/ in ELISA /ELISPOT diluent substituted with 20% mouse serum (in-house serum harvested from naïve C57BL/6JRj mice) and 50 µl were distributed per well. Plates were incubated for 2 h at RT and washed 3 times with PBS-T. Fifty μl per well of the biotinylated FLAMOD-specific detection Mab (4C1H7 mouse IgG1 - lot b220809-7034, [34]) diluted at 2 µg/ml in ELISA/ELISPOT diluent were distributed and incubated for 1 h at RT. Plates were washed 3 times with PBS-T and 50 μl per well of the avidin-HRP (eBioscience 18-4100-51) diluted 1:2000 in ELISA/ELISPOT diluent were added and incubated for 25 min at RT. Plates were washed 5 times with PBS-T. The reaction was developed using 50 μl per well of the TMB substrate reagent (Interchim ref: UP664781). Plates were incubated for 5 min at RT in the dark, prior to stopping the reaction by adding 25 μl H_2_SO_4_ 2N per well. The plates were read immediately at 450 nm absorbance and subtracted at 540-570 nm. The values were transformed using a four-parameter logistic regression (R^2^>0.99) to calculate the concentration of FLAMOD in the biological samples.

### 2.5 Transcriptional analysis by RT-qPCR and RNA sequencing

Total RNA was extracted with the Nucleospin RNA Plus kit (Macherey Nagel, Düren, Germany). Trachea and whole lungs were homogenized in RA1 buffer with an UltraTurrax homogenizer. Lysates of nasal cavity were obtained by injecting 500 μl RA1 buffer through a 22-gauge canula that was inserted into the posterior naris along the direction of the nostrils and collecting from the anterior naris. Bronchial epithelium sampling was performed as described [35]. Briefly, sanded PE50 tubing was inserted into the main bronchus via the trachea with gentle brushing and immediately processed for RNA extraction. Total RNA was reverse-transcribed with the High-Capacity cDNA Archive Kit (Applied Biosystems, Foster City, CA). The cDNA was amplified using SYBR green-based real-time PCR on a Quantstudio 12K PCR system (Applied Biosystems). Relative mRNA levels (2^−ΔΔCq^) were determined by comparing first the PCR cycle thresholds (Cq) for the gene of interest and the reference genes *Actb* and *B2m* (ΔCq), and then the ΔCq values for treated vs. control group (ΔΔCq). All the primers used in this study are listed in **Supplementary Table 1**.

For sequencing, RNA concentration and quality were evaluated by spectrophotometry (Nanodrop) and the TapeStation 4200 (Agilent Technologies), respectively. The libraries were prepared using the QIAseq stranded mRNA library kit (Qiagen), quantified by quantitative PCR (KAPA Library Quantification Kit for Illumina platforms; KapaBiosystems) and pooled in an equimolar manner prior to sequencing on a NovaSeq sequencer (Illumina) in 2 × 150 bp. RNA sequencing data (FASTQ files) were mapped to the mouse genome build (GRCm39) with STAR (STAR-2.7.10a). The mapped reads were annotated using GENCODE M30 GTF file. Differentially expressed genes analysis (DEG) were performed using the DESeq2 package (1.38.2 [36]). Median of ratio method was used for data normalization. Genes with null sum of read counts were filtered out. Genes with p-adjusted value < 0.05 and log2 fold change >1 were considered as differentially expressed. Over-Representation Analysis (ORA) was performed using ClusterProfiler (version 4.6.2) and the functions enrichGo and CompareCluster to identify Biological Processes. Analysis was carried out using Bioconductor (version 3.16.0). Sequencing data have been deposited in the National Center for Biotechnology Information’s Gene Expression Omnibus repository under the accession number GSE252037.

### 2.6 Cytokine and chemokine production

Protein concentrations of CCL20, CXCL2, IL-6, IL-22, and SAA were determined in BAL fluids and serum by enzyme-linked immunosorbent assay (ELISA kit from eBioscience, R&D Systems or Becton Dickinson).

### 2.7 Scintigraphy in mice

All scintigraphy experiments were conducted on male C57BL/6JRj (9-16 weeks old), approved by local ethics committee and recorded under the agreement reference N°11682-201700217166146. Prior to the scintigraphy study, a perfusion scintigraphy study was performed to evaluate technetium 99m (^99m^Tc) γ-rays absorption by tissues. Mice were slightly sedated via isoflurane inhalation (Vetflurane, Virbac, France) and a suspension of macro aggregates labeled with ^99m^Tc (^99m^Tc-MAA, Pulmocis, Curium Pharma, France) was intravenously injected through the tail vein. After ensuring that all ^99m^Tc-MAA was trapped in the pulmonary capillaries, animals were imaged in anterior static position using Biospace® gamma imager (Biospace, France) for 60 seconds under isoflurane anesthesia. This step allowed to determine the attenuation coefficient of animal tissues against γ-rays, which must be considered when calculating activity measured in mice, and takes into account the background activity and radioactive decay of technetium. Moreover, it allows the illustration of the region of interest (ROI). For the scintigraphy study, inhalation by intranasal and nebulization routes were compared. For intranasal delivery mice were sedated, then 40 µL of 0.9% NaCl solution mixed with diethylenetriaminepentaacetic acid labeled with ^99m^Tc (^99m^TC-DTPA, Technescan™ DTPA, Curium Pharma, France) were rapidly administered. For the nebulization route, experiments were performed as described in 2.3 using the custom-made nebulization chamber and an Aerogen^®^ Solo to nebulize 1 mL of 0.9% NaCl solution mixed with 0.1 mL of ^99m^TC-DTPA (n=4 per nebulization round). Immediately after i.n. or nebulized administration, mice were imaged for 120 seconds under isoflurane anesthesia.

### 2.8 Histology and staining

The left lung from each mouse was collected in buffered 10% formalin (Q-Path VWR Chemicals), included in paraffin, and 3- to 5-µm tissue sections were prepared (Althisia, Troyes, France). Tissue sections (n=3 for FLAMOD treated and n=2 for untreated control) were stained with a classical hematoxylin and eosin reagent or processed for immunohistochemistry. Immunohistochemistry on neutrophils was performed using rabbit monoclonal anti-Ly6G antibody (clone EPR22909-135). The Ly6G-stained images were evaluated for neutrophil infiltration.

### 2.9 Statistical Analysis

Results were expressed as individual values and median [interquartile range] or mean ± standard error of the mean (SEM). Groups were compared using a Kruskal-Wallis one-way analysis of variance (ANOVA) with Dunn’s post-test (for three or more groups) and two-way ANOVA mixed model with Šídák’s multiple comparisons test. All analyses were performed using GraphPad Prism software (version 10.10, GraphPad Software Inc., San Diego, CA), and the threshold for statistical significance was set to p<0.05.

## 3. Results

### 3.1 Fate of inhaled FLAMOD in mice after intranasal and aerosol administration

Understanding the fate of inhaled flagellin is imperative to characterizing its safety and efficacy parameters as an adjunct to antibiotics. The outcome of flagellin (defined here as FLAMOD) was investigated after intranasal administration of 30 μg FLAMOD in bronchoalveolar lavages (BAL), as a surrogate of lung deposition, and serum using a FLAMOD-specific ELISA. Immediate recovery after administration (<5 min) revealed that around 30 % of the administered FLAMOD reached the lungs, *i.e.,* 8.6 ± 2.0 µg (**Figure 1A**). Similar levels of lung deposition (∼45 %) were defined using scintigraphy (**Figure 1C-D**). FLAMOD was rapidly degraded over time and became undetectable after 24 h. In addition, serum levels of FLAMOD remained consistently below the lower limit of quantification (LLOQ), *i.e.,* <4 ng/mL, at all timepoints following administration.

**Figure 1.**
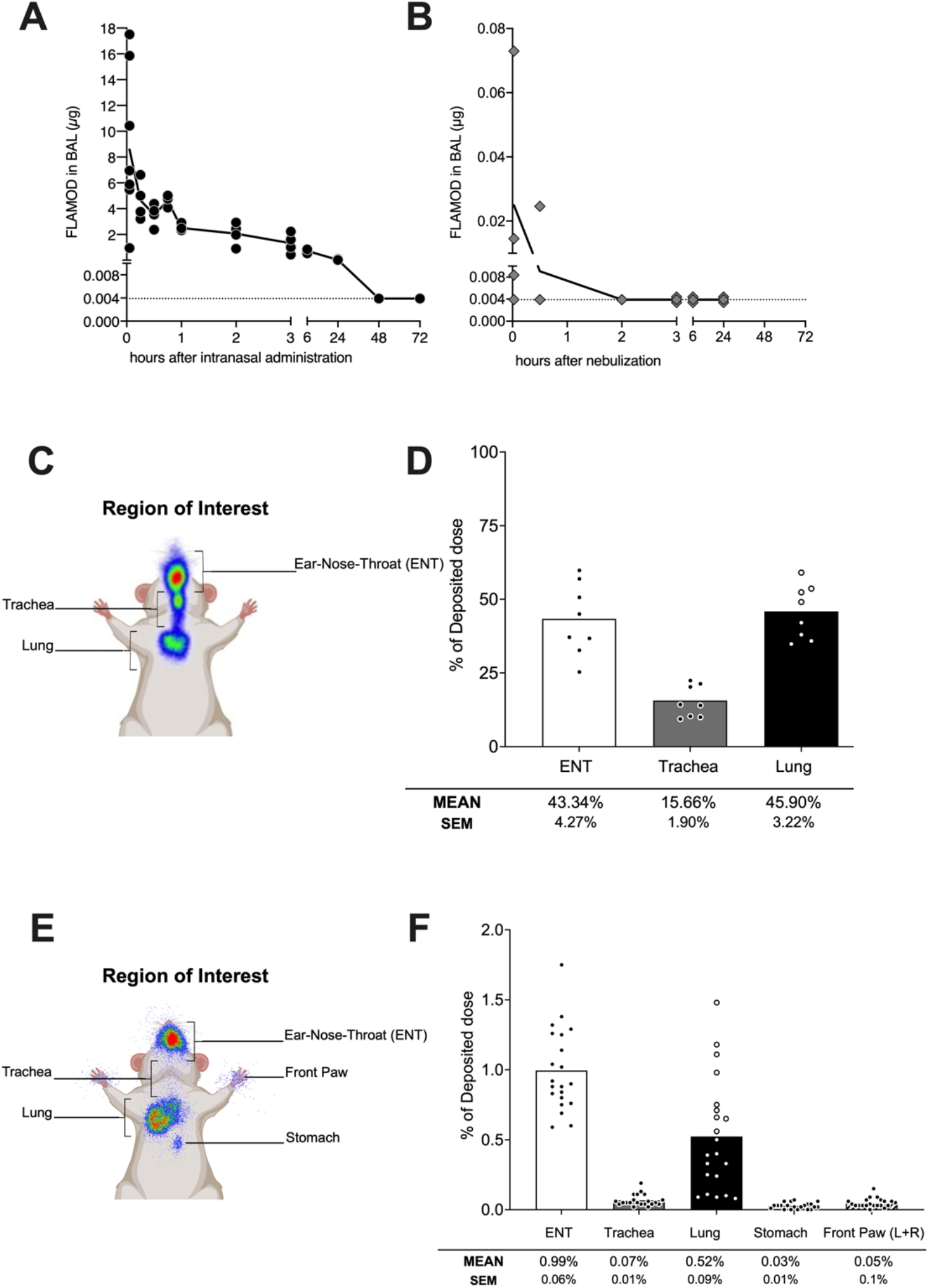
FLAMOD biodistribution after intranasal or aerosol inhalation. BALB/c mice (n=4-8) per time point and treatment group) were subjected to intranasal administration of 30 μg FLAMOD or nebulization with 1 ml of FLAMOD at 250 μg/ml in NaPi buffer, utilizing a mesh-nebulizer and aerosol chamber setup. Control groups were either treated with NaPi buffer or left untreated. BAL was collected at varying timepoints to measure FLAMOD. **(A-B)** FLAMOD deposition in BAL after **(A)** intranasal administration or **(B)** nebulization was quantified using a FLAMOD-specific ELISA. Dotted line indicates LLOQ of FLAMOD, i.e., 3.9 ng/ml. Deposition of intranasal FLAMOD (**C-D**) and nebulized FLAMOD (**E-F**) was further analyzed by scintigraphy. Mice were subjected to intranasal administration of 0.9% NaCl containing ^99m^TC-DTPA (n=8), or nebulization with 0.9% NaCl containing ^99m^TC-DTPA (n=21). Left images show the regions of interest (ROI) where radioactivity was measured, and the quantitative data for each ROI are presented on the right panel. The amount of aerosol deposited is expressed as a percentage of the total amount administered.

Mice were also exposed to nebulized FLAMOD (1 ml at 250 µg/mL) for 5 min using a customized chamber connected to an Aerogen® Solo mesh nebulizer (**Supplementary Figure 1A-B**). FLAMOD deposition in BAL reached 25 ± 16 ng immediately after aerosolization and decreased to below the LLOQ within the first 2 h (**Figure 1B**). In contrast, scintigraphy determined that 0.52 % lung deposition after nebulization is achieved which is equivalent to 1.30 µg FLAMOD (**Figure 1E-F**). FLAMOD was undetectable in serum post-nebulization, indicating no systemic leakage. The difference in lung deposition between the two analytical methods (ELISA and scintigraphy) highlights differing sensitivities and suggests rapid degradation of FLAMOD upon aerosol delivery. To better understand the fate of FLAMOD when in the circulatory compartment and to have a positive reference for immune activation when FLAMOD is found directly in the systemic compartment, FLAMOD was injected intravenously (**Supplementary Figure 2**). Serum levels of FLAMOD corresponding to around 60% delivered dose were detected immediately after injection (< 5 min) followed by a decline to <1% by 60 min post-administration. This suggests rapid FLAMOD clearance. In conclusion, inhalation of FLAMOD by the intranasal or nebulization route in mice provides local delivery without systemic leakage, while resulting in distinct levels of lung deposition.

### 3.2 Immunostimulatory activity of inhaled FLAMOD

We first confirmed the TLR5-dependent activation of immune mediators by FLAMOD using TLR5 knock-out mice. As expected, when FLAMOD was provided intravenously to wild-type and TLR5 knock-out mice respectively, no immune mediators (namely CCL20, IL-6 and IL-22) were produced in the TLR5 knock-out mice in response to FLAMOD treatment (**Supplementary Figure 3**). We next analyzed the TLR5-specific immune responses to nebulized FLAMOD based on the expected immune outcomes after intranasal delivery [11, 32, 37–39]. The activation of respiratory innate immunity subsequent to intranasal or nebulization delivery of FLAMOD exhibited comparable levels of CCL20, CXCL2 and IL-6 in BAL (**Figure 2A**). However, aerosol delivery resulted in a less persistent immune response compared to intranasal administration, generally returning to steady-state within a shorter period to that of intranasal. Serum analysis revealed that nebulization triggered similar levels of IL-6 and CCL20, as well as shorter responses compared to intranasal delivery (**Figure 2B**). The serum concentration of the acute phase inflammation marker serum amyloid A (SAA) was globally lower in response to nebulization than intranasal delivery. Importantly, IL-22 was undetectable in serum after nebulization in contrast to intranasal route. We previously showed that IL-22 is produced when flagellin is found systemically but not when it stays in the mucosal compartment [39]. Thus, this cytokine indicates leakage of flagellin into the bloodstream and thus represents a surrogate marker of lung barrier integrity. The absence of IL-22 in the serum of nebulized mice suggests that FLAMOD does not cross the pulmonary barrier after nebulization, particularly when compared to the very high IL-22 levels detected in mice receiving FLAMOD intravenously (**Supplementary Figure 2**). These findings fully corroborate the data on FLAMOD biodistribution presented above.

**Figure 2.**
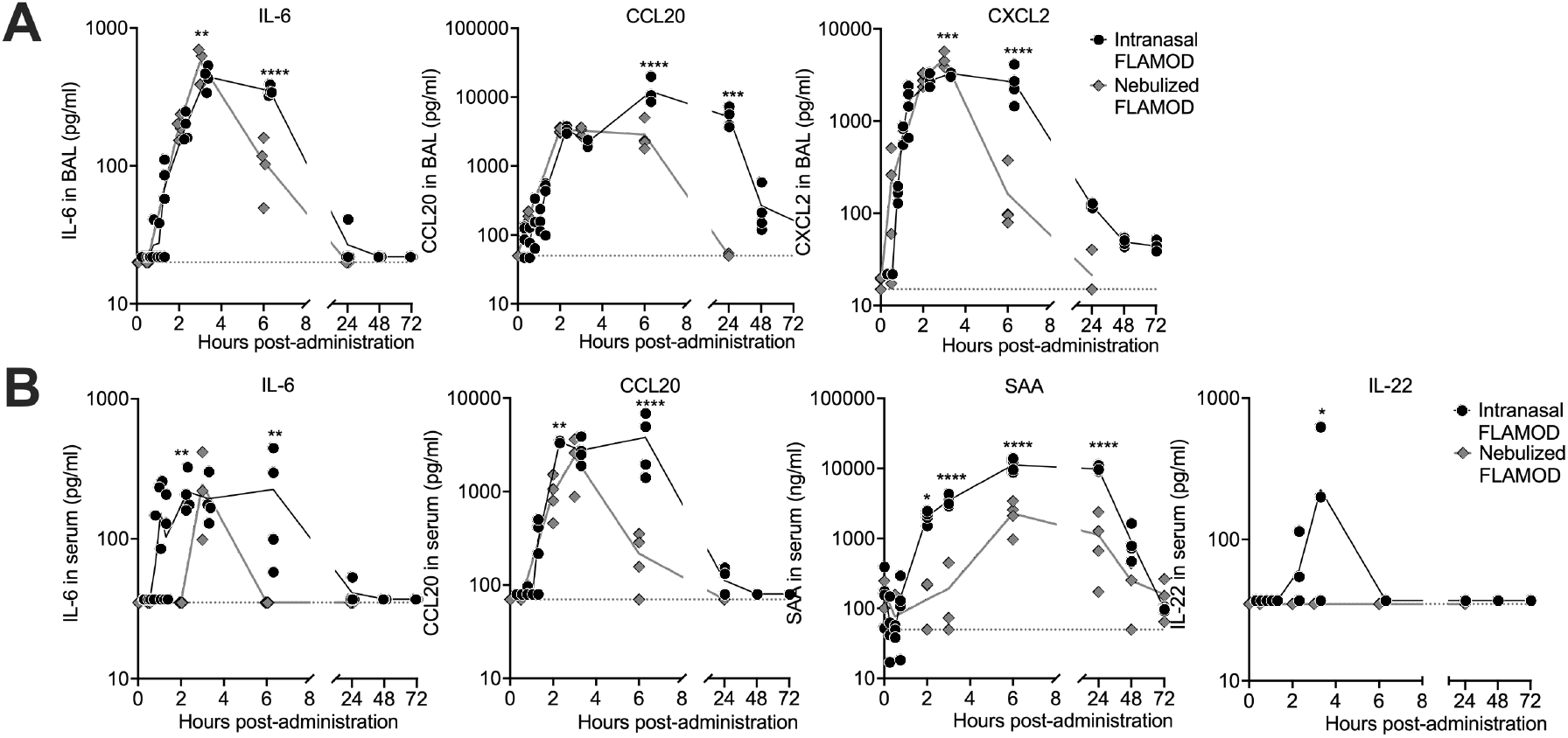
Transient innate immune activation following FLAMOD inhalation. BALB/c mice (n=4 per time point and treatment group) were subjected to intranasal administration of 30 μg FLAMOD or nebulization with 1 ml of FLAMOD at 250 μg/ml in NaPi buffer, utilizing a mesh-nebulizer and aerosol chamber setup, or left untreated. BAL and serum were collected at varying timepoints to monitor immune response. Cytokine and chemokine concentrations in BAL fluids **(A)** and serum **(B)**. Dotted lines indicate LLOQ. Statistical differences were analyzed using two-way ANOVA (*p<0.05, **p<0.01, ***p<0.001, ****p<0.0001).

Consistent with the findings from ELISA analysis of BAL (**Figure 1**) and similarly to the previous finding following intranasal delivery [11, 32, 37], nebulized FLAMOD elicited a temporary increase in transcriptional activity of pro-inflammatory genes, reaching its peak at 2 h post-administration (**Supplementary Figure 4)**. Given that inhalation delivers the drug into the nose, trachea, bronchi and lung, we compared the gene expression from these compartments 2 h after the treatment of animals with nebulized or intranasal FLAMOD and normalized the expression profiles to the buffer treated animals (**Figure 3**). Inhaled FLAMOD stimulated the transcription of *Il6*, *Ccl20* and *Cxcl2* in all respiratory compartments. Interestingly, with the exception of the nasal compartment, activation levels were significantly higher under the nebulization condition than for intranasal inhalation. This finding is also intriguing when considering that scintigraphy data displayed negligible tracheal deposition during nebulization (0.07%) as opposed to that deposited intranasally (15.7%), amounting to a 224-fold difference. To further compare the modes of inhalation, RNA-seq analysis was conducted on lungs collected at 2 h after FLAMOD or buffer administration (**Figure 4 and Supplementary Table 2**). This analysis identified 955 and 1470 differentially expressed genes (DEG) that were significantly regulated (log_2_FC≥1 and p≤0.05) by aerosol and intranasal inhalation, respectively. Similar genes were up- and down-regulated for both types of FLAMOD administration as depicted in **Figure 4A-B and Supplementary Table 2**. For both inhalation modes, over-representation analysis (ORA) revealed a significant modulation of signaling pathways associated with response to bacterial molecules, innate immunity and innate receptors, NF-κB, cytokine/chemokine production, immune cell activation and migration, defense to pathogens or negative regulation to dampen inflammation (**Figure 4C and Supplementary Table 3**). Despite the significant disparity in FLAMOD deposition observed across nebulization and intranasal delivery, our findings demonstrated that nebulization is a potent method for eliciting TLR5-specific innate immune responses within the respiratory tract.

**Figure 3.**
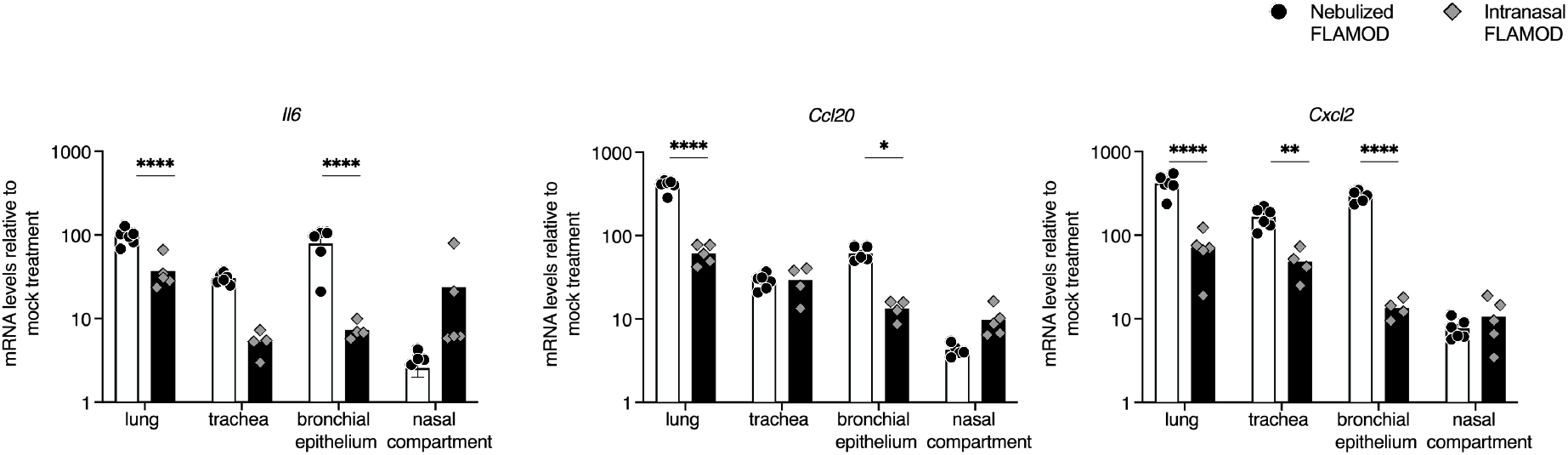
Compartmentalization of transcriptional response to FLAMOD inhalation. BALB/c mice (n=5-6 per time point and treatment group) were subjected to intranasal administration of 30 μg FLAMOD or nebulization with 1 ml of FLAMOD at 250 μg/ml in NaPi buffer, utilizing a mesh-nebulizer and aerosol chamber setup, or mock treatment with NaPi buffer. Lungs, trachea, bronchial epithelium and nasal lavages were analyzed at 2 h post-administration. Gene expression of innate immune markers was measured by RT-qPCR. The relative mRNA levels of selected genes are expressed against the reference gene *Actb* and normalized to the mock condition. Statistical differences were analyzed using two-way ANOVA (*p<0.05, **p<0.01, ****p<0.0001).

**Figure 4.**
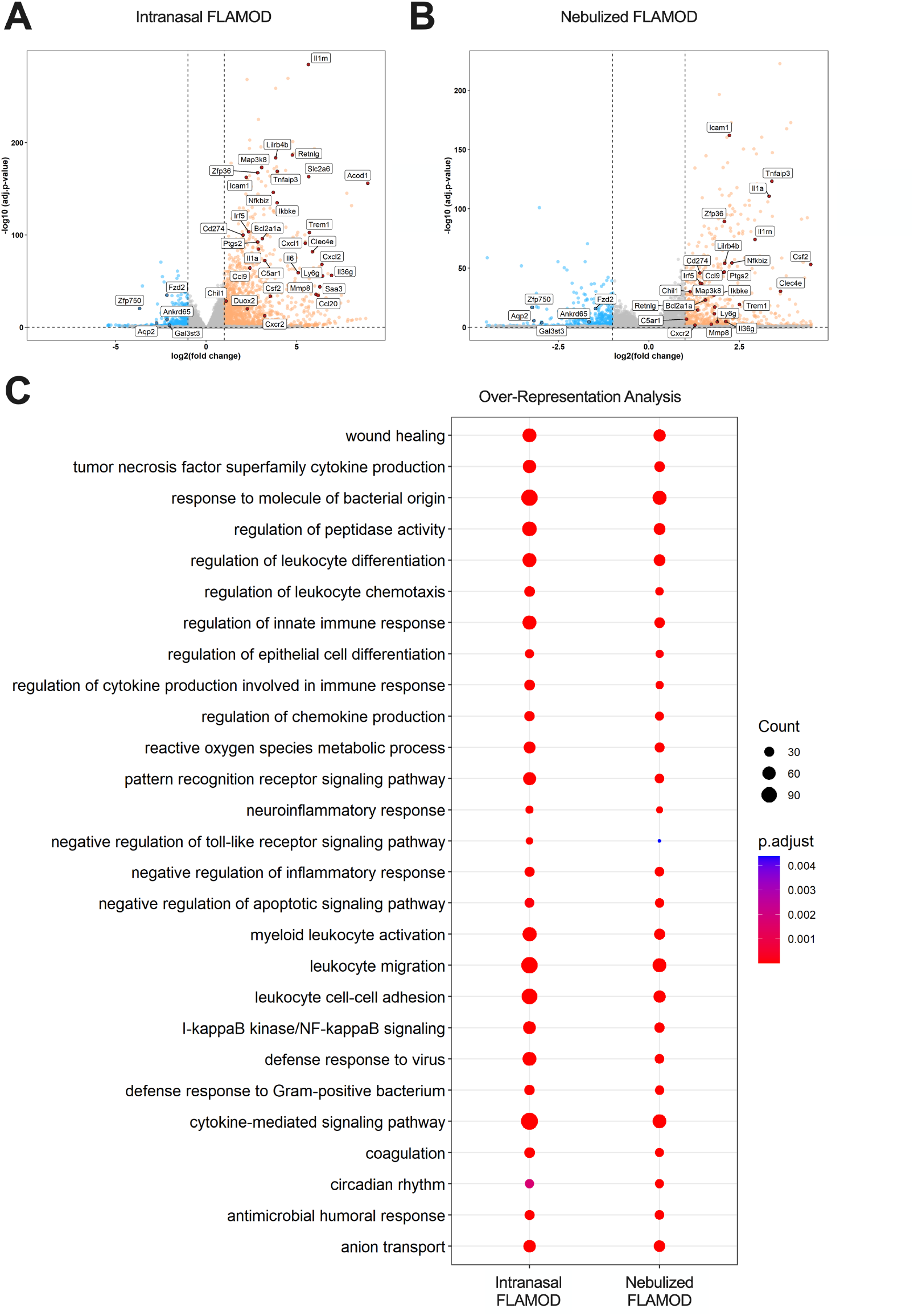
FLAMOD inhalation by the intranasal or nebulization route activates similar innate immune responses. BALB/c mice (n=4 per time point and treatment group) were subjected to intranasal administration of 30 μg FLAMOD or nebulization with 1 ml of FLAMOD at 250 μg/ml in NaPi buffer, utilizing a mesh-nebulizer and aerosol chamber setup, or mock treatment with NaPi buffer. Lungs were collected 2 h post-administration for RNA sequencing. **(A-B)** Volcano plots of DEG between intranasal FLAMOD and intranasal buffer group **(A)** and nebulized FLAMOD and nebulized buffer group **(B)**. Labelled genes represent the top DEG. **(C)** Immune pathways in nebulized and intranasal groups were identified using ORA. The common biological processes enriched in both conditions are represented. The count and p.adjust scales indicate the number of genes enriched in the pathway and the significance of enrichment, respectively.

Neutrophil recruitment stands as a pivotal indicator of TLR5 activation within pulmonary tissues [10, 16, 37, 40, 41]. To elucidate the influence of inhalation on this process, lung samples were analyzed by immunohistochemical staining employing the anti-Ly6G antibody for neutrophil detection (**Figure 5**). Evaluations were performed at two distinct time points: 4 hours post-treatment, coinciding with the peak of neutrophil recruitment in response to flagellin [10, 37], and 24 hours post-treatment, representing a period wherein FLAMOD becomes undetectable and most activation markers return to baseline levels. After 4 h, a comparable pattern of neutrophil infiltration was observed between the two inhalation methods, with a tendency for more clustering around the bronchioles rather than within the alveolar space. However, a notable contrast emerged at 24 h between the intranasal and nebulized conditions. In the intranasal group neutrophil presence remained elevated, while in the nebulized group the level of infiltration resembled that of untreated controls. Taken together, these observations suggest that nebulized FLAMOD can induce transient lung neutrophil infiltration, with a more rapid return to baseline compared to intranasal administration.

**Figure 5.**
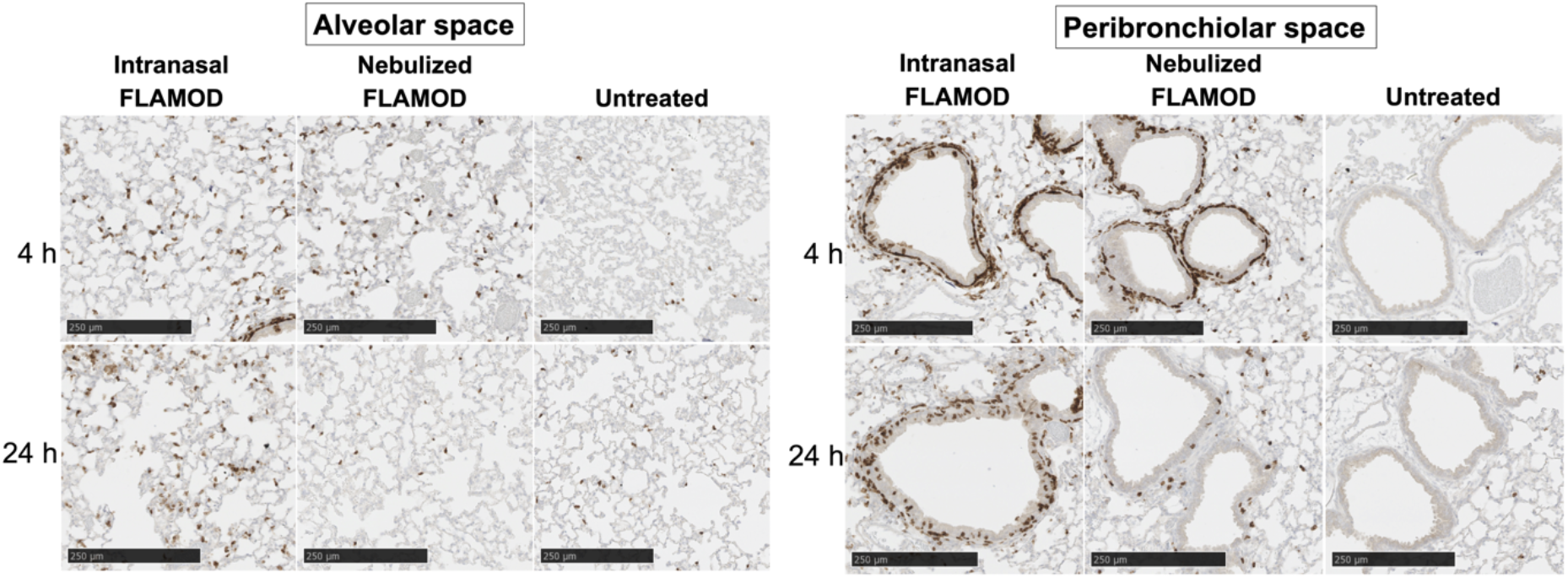
Transient neutrophil recruitment in response to FLAMOD inhalation. BALB/c mice (n=3 per FLAMOD treatment) were subjected to intranasal administration of 30 μg FLAMOD or nebulization with 1 ml of FLAMOD at 250 μg/ml in NaPi buffer, utilizing a mesh-nebulizer and aerosol chamber setup, or mock treatment with buffer (n=2). Lungs were collected 4 h and 24 h after FLAMOD inhalation and processed for immunohistochemical Ly6G-staining of neutrophils (brown). (**A)** Alveolar space and **(B)** peribronchiolar space. The images are representative of individual animals.

### 3.3 Therapeutic efficacy of nebulized FLAMOD against pneumococcal pneumonia

Nebulization of FLAMOD was next compared to intranasal delivery of FLAMOD as an adjunct therapy to the antibiotic amoxicillin (AMX) in a pneumococcal superinfection model (**Figure 6A**). To this end, the animals were superinfected with a serotype 19F of *S. pneumoniae* that displays resistance to AMX (MIC_AMX_, of 1.5 μg/ml). As expected, treatment with AMX and intranasal FLAMOD substantially improved bacterial clearance compared to AMX alone (**Figure 6B**). The same synergy and efficacy were found when FLAMOD was nebulized resulting in significant reduction of bacterial burden in the lung as well as bacterial dissemination in spleen (**Figure 6C**). In the context of superinfection, the fate of FLAMOD in the BAL after intranasal inhalation or nebulization was in line with that observed for naïve mice (**Figure 6D-E**). However, FLAMOD was found at very low levels in the bloodstream of mice treated intranasally, suggesting marginal leakage (**Figure 6D**). The immune activation profiles in lung and bloodstream were also similar to those seen in mice without prior infection, implying that activity of inhaled FLAMOD did not change in the context of infection (**Figure 6F-G**). Importantly, IL-22 was not detected in the bloodstream after nebulization with FLAMOD, indicating that even when the lungs are compromised by infection, leakage is not observed. Finally, the therapeutic synergy of nebulized FLAMOD with AMX was also effective in a primary respiratory infection with a serotype 1 amoxicillin-sensitive strain (MIC_AMX_ = 0.016μg/ml) (**Supplementary Figure 5**). Collectively, these data indicate that despite the low lung deposition upon nebulization in mouse models of antibiotic-resistant and antibiotic-sensitive pneumococcal infections, the delivered dose is sufficient to provide therapeutic efficacy within the range observed for intranasal administration.

**Figure 6.**
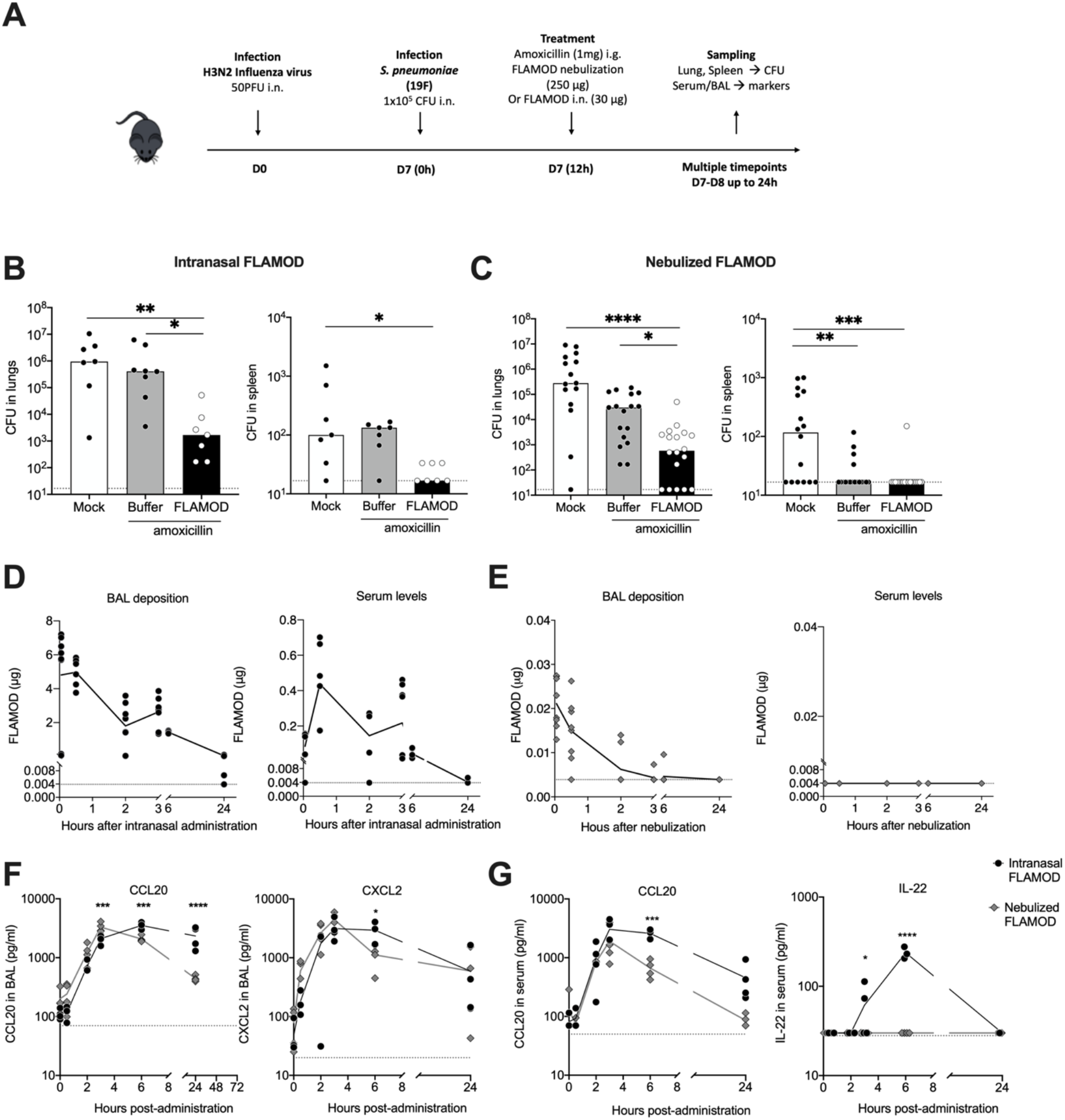
FLAMOD inhalation efficacy, kinetics and therapeutic activity in pneumococcal superinfection. **(A)** C57BL/6 mice (n=7-8 for intranasal and 15-19 for nebulization, per condition) were infected intranasally with influenza virus followed 7 days later by intranasal inoculation of 1×10^5^ CFU of *S. pneumoniae* 19F. Animals were treated at 12 hours post pneumococcal infection by intranasal administration of 2.5 µg **(B, F-G) or** 30 μg **(D)** FLAMOD or nebulization with 1 ml of FLAMOD at 250 μg/ml in NaPi buffer **(C, E-G)** combined with intragastric administration of 1 mg amoxicillin. Mock animals received NaPi buffer and water. **(B-C)** Lungs and spleen for CFU were sampled 12 h post-treatment to monitor therapeutic effects of intranasal administration **(B)** or nebulized FLAMOD **(C)**. Statistical differences between the groups were analyzed by One-way ANOVA (*p<0.05, **p<0.01, ***p<0.001, ****p<0.0001). **(D-G)** BAL and serum were sampled to measure FLAMOD and immune activity. **(D-E)** FLAMOD distribution in BAL and serum after intranasal administration **(D)** and nebulized FLAMOD **(E)**. FLAMOD was measured by ELISA. LLOQ was fixed to 4 ng/ml. **(F-G)** Cytokine and chemokine production in BAL **(F)** and serum **(G)** in response to FLAMOD inhalation methods. Statistical differences between the two methods over time were analyzed using two-way ANOVA (*p<0.05, **p<0.01, ***p<0.001, ****p<0.0001).

## 4. Discussion

This study underscores the potent immunostimulatory effects of nebulized flagellin (FLAMOD) on the respiratory tract’s immune defenses by combining quantitative analysis of the biologic and the immune responses to it in the airways and bloodstream, as well as its therapeutic efficacy. Although FLAMOD has been administered intranasally to target the lower respiratory tract in mice, this route is not appropriate for similar targeting in humans. To overcome this limitation and bridge the translational gap for this therapeutic, we engineered a new custom-designed chamber integrated with the Aerogen Solo^®^ vibrating mesh nebulizer. Despite observing low lung deposition of nebulized FLAMOD, our results indicate that the immunomodulatory and therapeutic efficacy of FLAMOD remained potent. Remarkably, our aerosol delivery system achieved comparable therapeutic outcomes to intranasal administration against pneumococcal pneumonia [15, 16], despite delivering doses 5-to-300-fold lower, as defined by scintigraphy and ELISA data, respectively. Furthermore, nebulization was not linked to any significant systemic exposure at the deposited dose achieved, thereby highlighting both the relevance of the delivery methodology and the safety profile. Our findings provide robust evidence supporting the development of FLAMOD as an immune-booster complementing antibiotic therapy against respiratory bacterial pathogens with high potency even at low concentrations.

Scintigraphy analysis unveils distinct contrast in deposition patterns between nebulized and intranasally administered FLAMOD. A small fraction, less than 1%, of the nebulized formulation is found to reach the lower airways, contrasting sharply with the intranasal route, which achieves a deposition exceeding 40%. Furthermore, scintigraphy demonstrates pronounced divergence in distribution across respiratory compartments between nebulized and intranasal routes. Notably, the intranasal approach exhibits heightened exposure of the trachea, whereas such exposure is lacking in the nebulization method. Despite these differences in deposition patterns, RNAseq analysis indicates activation of similar immune pathways in the lungs across both administration approaches. Moreover, RT-qPCR analysis of genes encoding immune mediators reveals comparable or even elevated levels of transcriptional activation in the nasal compartment, trachea, and bronchial epithelium following nebulization, in contrast to intranasal administration. These findings suggest that saturation of immune activation is achieved at low doses of FLAMOD effect. While it has been previously inferred that inhalation of flagellin does not enter the bloodstream [11], this study was the first to measure FLAMOD using a new ELISA test developed for this purpose. Compared to scintigraphy, ELISA measurements reported much lower levels in the deposition of FLAMOD in the pulmonary compartment shortly after administration: less than 0.1% for nebulized FLAMOD versus 30% for intranasal administration. These findings strongly suggest rapid elimination of nebulized FLAMOD within the airways, despite the differential deposition detection by the two quantification methods used.

Several factors could contribute to accelerated FLAMOD elimination upon nebulization. The intranasal administration facilitates the entry of the biologic solution into the airways primarily though gravitational forces and diffusion mechanisms. In contrast, nebulization generates fine liquid particles that deeply penetrate tissues via the respiratory flow [42]. The interaction of nebulized particles with various components such as epithelial lining fluid factors, mucus, pulmonary surfactants, epithelial cells and alveolar macrophages is expected to be of a specific nature. Consequently, these distinct inhalation methods lead to differing spatial distributions and interactions within the respiratory system, which may play a crucial role in the observed degradation dynamics of FLAMOD. Previous studies have reported that proteins deposited in the airways by nebulization can be eliminated by various mechanisms including pulmonary surfactant opsonization [43], endocytosis or phagocytosis by alveolar macrophages [44] proteolytic degradation, mucus entrapping, or mucociliary clearance [45]. It is usually considered that mucus is not a barrier for proteins <500 kDa while particles trapped in the mucus, can be transported to the pharynx by the mucociliary clearance mechanisms [45, 46]. The conducting airways including bronchi and alveoli contain many secreted and cell-associated proteases such cathepsins, dipeptidylpeptidases, metalloproteinases, carboxypeptidases, or xenobiotic metabolizing enzymes such as cytochromes that can be active against biologics [47, 48]. Of note, FLAMOD is derived from the native *Salmonella* flagellin that is known to contain multiple proteolytic cleavage sites in the distal region, rendering FLAMOD prone to degradation by proteases [49]. Whether nebulization exacerbates these catabolic processes compared to intranasal delivery remains to be determined.

Biologic drugs specific for treating lung conditions, such as the recombinant human DNAse used as a mucolytic in cystic fibrosis lung disease, have been adapted for nebulization as it is a unique method to compartmentalize the drug activity within the respiratory tract, thereby mitigating systemic exposure and potential toxicity [50, 51]. While the TLR5-dependent immunomodulation in the airways presents a promising therapeutic avenue, it is important to acknowledge a potential caveat stemming from systemic exposure. TLR5, being widely expressed across various cells and tissues, carries the risk of precipitating undesired inflammation, commonly referred to as immune-related adverse events [52]. Nebulization is thus an attractive administration route for our biologic in order to mitigate such events. To assess systemic exposure to FLAMOD, the production of serum IL-22 serves as a surrogate marker. Previous studies have established that IL-22 necessitates flagellin-mediated activation of TLR5 signaling in dendritic cells derived from either blood or tissue parenchyma [11, 38, 39]. This study showed no or very low levels of serum IL-22 after inhalation, corroborating the FLAMOD dosing in serum and insinuating little to no systemic leakage. The efficient elimination of nebulized FLAMOD may contribute to the minimal systemic exposure observed. Factors that influence systemic absorption include properties of the aerosol particles such as pH, charge, and surface activity. Studies have indicated a variable availability of inhaled proteins in the bloodstream, ranging from 10% to 50% [53]. In mice, systemic exposure to antibodies via orotracheal inhalation is notably low, despite efficient deposition in the airways [54]. This trend is echoed in clinical trials, where nebulized administration of antibodies results in systemic exposure of approximately 1% of the deposited dose [55].

Despite the rapid clearance of FLAMOD, its administration via nebulization still effectively stimulates innate immunity within the lungs, mirroring the response observed with the intranasal route. TLR signaling, including activation of TLR5, initiates immediate NF-κB activation within minutes, alongside the induction of negative feedback mechanisms [8, 9]. This “hit and run” activation of TLR5 rationalizes the efficacy of nebulized FLAMOD despite its short residency half-life within respiratory tissues. Although nebulized FLAMOD induces a transient immunomodulatory effect on neutrophil mobilization in alveoli and bronchi, it promotes a more rapid resolution of the TLR5-mediated pro-inflammatory response compared to intranasal delivery. This accelerated resolution aligns with the observed differences in the biodistribution of intranasal versus nebulized FLAMOD, with intranasal FLAMOD persisting longer in the lungs compared to the nebulized form. However, the precise mechanisms underlying these differential kinetics require further investigation.

The effectiveness of nebulized FLAMOD as an adjunct to antibiotics was demonstrated in two clinically relevant models of pneumonia: primary pneumonia and pneumococcal superinfection, which occurs when there is a pre-existing influenza virus infection. Furthermore, this proof-of-concept was confirmed using two different serotypes of pneumococcus, one susceptible and the other resistant to amoxicillin (with MIC_AMX_ values of 0.016 μg/ml and 1.5 μg/ml respectively). The results from this study align with the clinical development of nebulized FLAMOD and support previous findings from intranasal administration in pneumonia treatment [15, 16]. Analysis of FLAMOD distribution throughout the body, as well as the immune responses in the lungs and blood following inhalation, revealed that pneumonia-induced damage had minimal impact on the observed bioavailability and immune activation patterns. This reinforces the safety and effectiveness of our therapeutic approach targeting the respiratory tissues.

In conclusion, our findings underscore the intricate interplay between inhalation methods, respiratory and systemic distribution, and immunomodulatory activity of FLAMOD, which are critical considerations in optimizing therapeutic outcomes and minimizing adverse effects. It will be important to extend these investigations to the study of more representative models to allow better translation of nebulized FLAMOD to the clinic. More appropriate species, for better defining nebulized FLAMOD PK and PD parameters with higher translational value, will include pig models and non-human primate models. Nevertheless, this study has determined nebulization as an effective and safe method for next step development of host-directed lung-targeting therapies of pneumonia.

## Author Contributions

MB, CCo, JP planned studies, performed experiments and analyzed data. DC, DS performed experiments. YZ analyzed transcriptomic data. RM provided the nebulizers plus controllers and reviewed the paper. FW provided the clinical isolate LILPNEUHC 19F strain and reviewed the paper. MB, CCo, YZ, JCS and CCa wrote the paper with input from all authors. MB, JCS, CCa and NHV supervised the project.

## Acknowledgements

We thank the Vaccine Development department of Statens Serum Institut (Denmark) for the manufacture and supply of FLAMOD. We thank the PLETHA animal facility from UMS2014-US41-PLBS.

