## Supplementary Figures_1_5 for "Targeted delivery of flagellin by nebulization offers optimized respiratory immunity and defense against pneumococcal pneumonia"

**The supplementary materials include:**

- **5 supplementary figures (attached to this document)**
- **3 supplementary tables (accessible as individual files)**

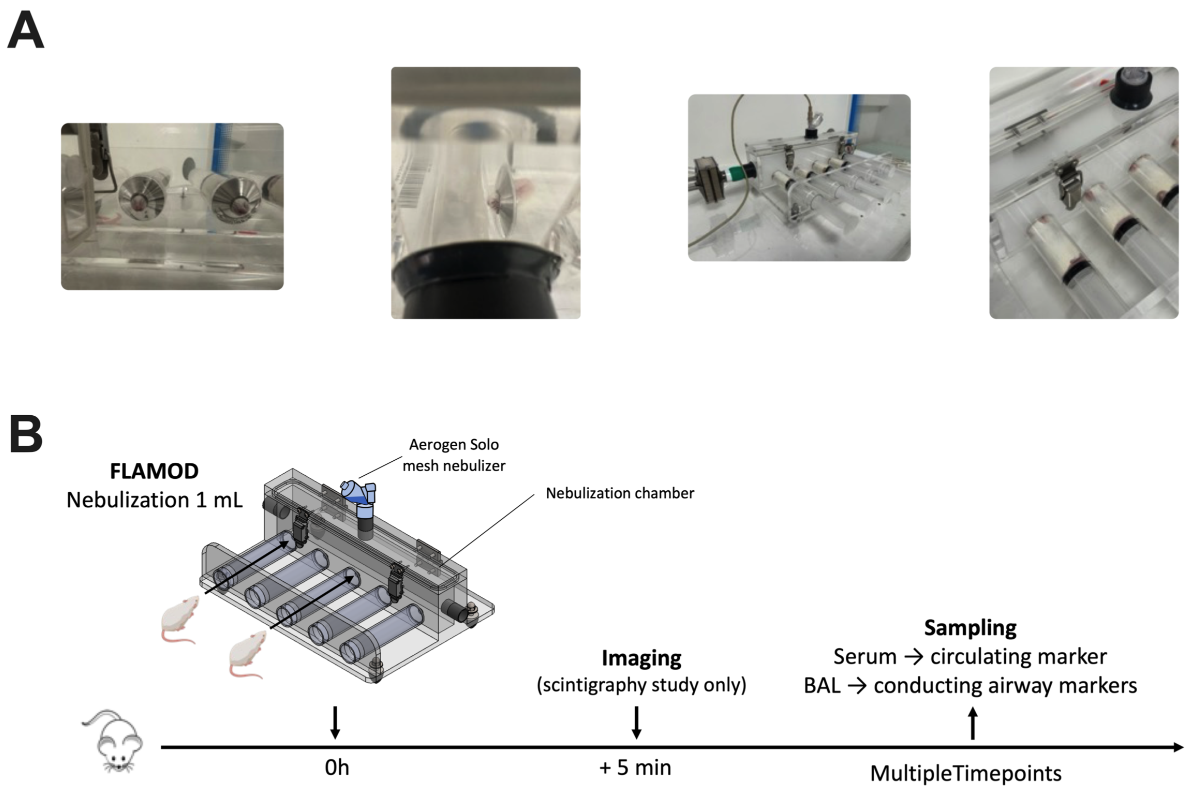


**Supplementary Figure 1. Custom-made nebulization chamber and experimental designs.** (**A**) Nebulization chamber images. Left images depict the naris of mice being exposed to the chamber where inhalation takes place. Right images describe the nebulization chamber set-up, including the Aerogen Solo vibrating mesh nebulizer. **(B)** Experimental design for kinetics and activity of nebulized FLAMOD in naïve mice.

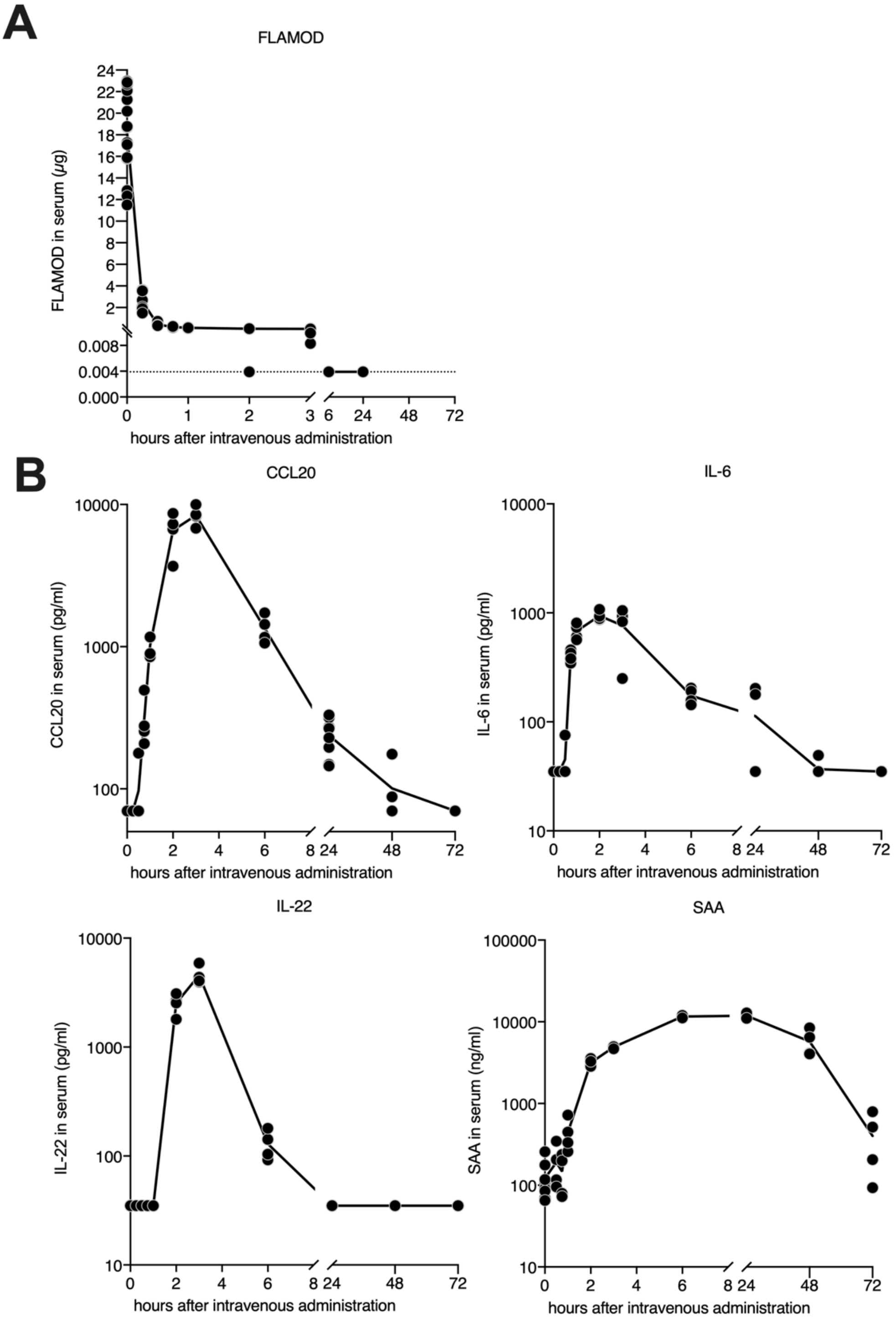


**Supplementary Figure 2. FLAMOD kinetics and activity upon intravenous administration.** BALB/c mice (n=4 per timepoint) were injected intravenously with 30 µg of FLAMOD (in 200 µl PBS). Serum was collected at varying timepoints to measure FLAMOD **(A)** and immune activity **(B)** FLAMOD, cytokines, chemokines and inflammatory markers were quantified by ELISA.

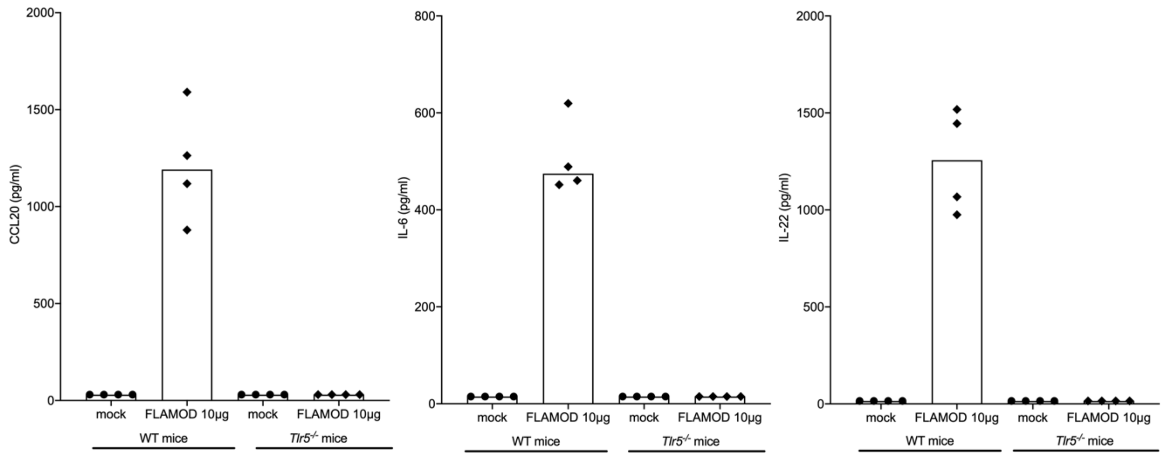


**Supplementary Figure 3. TLR5-specific activation by FLAMOD.** C57BL/6 mice (wild-type (WT) or *Tlr5*$^{-/-}$, n=4 per group) were injected intravenously with 10 µg FLAMOD in 200 µl PBS. Mock animals received 200 µl PBS alone. Blood was collected 2-hours post-treatment and TLR5 activation was assessed by measuring the serum concentrations of CCL20, IL-6 and IL-22 by ELISA.

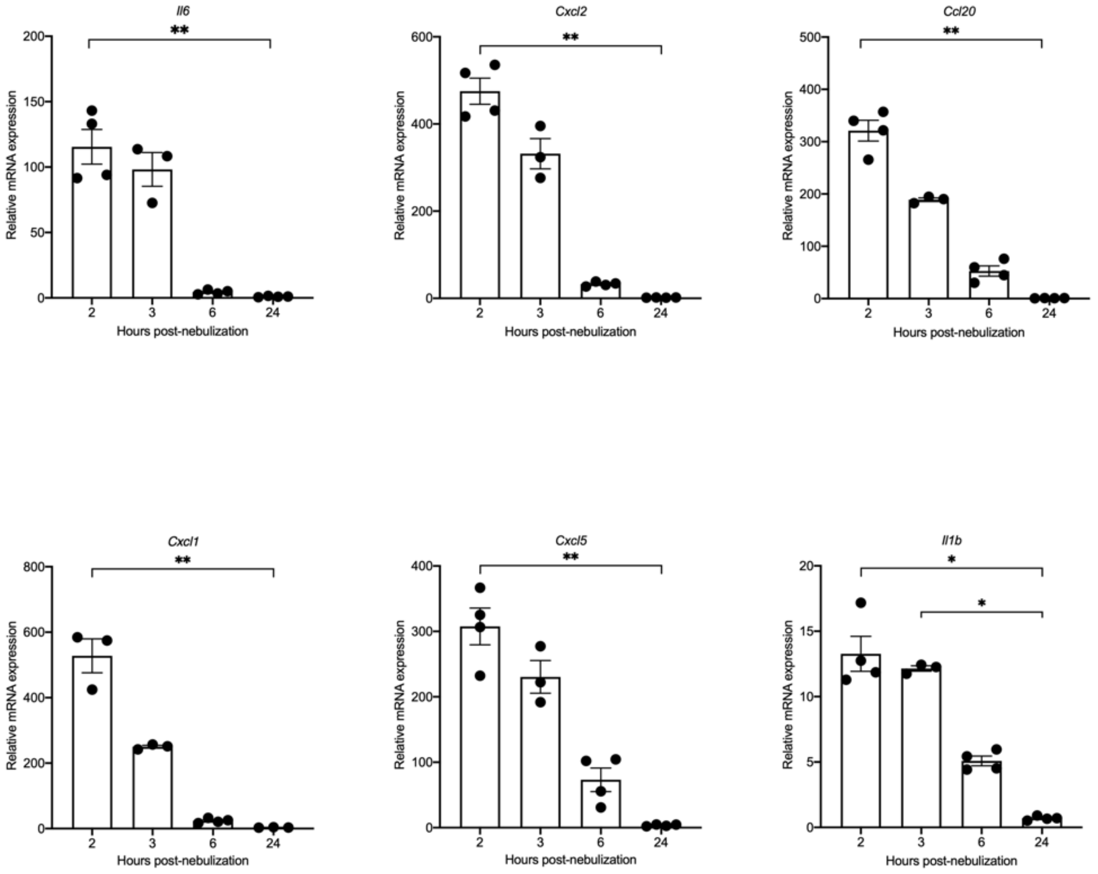


**Supplementary Figure 4. Time course of nebulized FLAMOD-mediated transcriptional activation in lungs.** BALB/c mice (n=4 per time point and treatment group) were treated, utilizing a mesh-nebulizer and aerosol chamber setup, with 1 ml of FLAMOD at 250 µg/ml in NaPi buffer or mock-treated with NaPi buffer as a control. Lungs were collected at varying timepoints to monitor expression of pro-inflammatory genes. The relative mRNA levels are expressed against the reference gene *Actb* and normalized to the mock condition. Statistical differences were analyzed using one-way ANOVA (*p<0.05, **p<0.01).

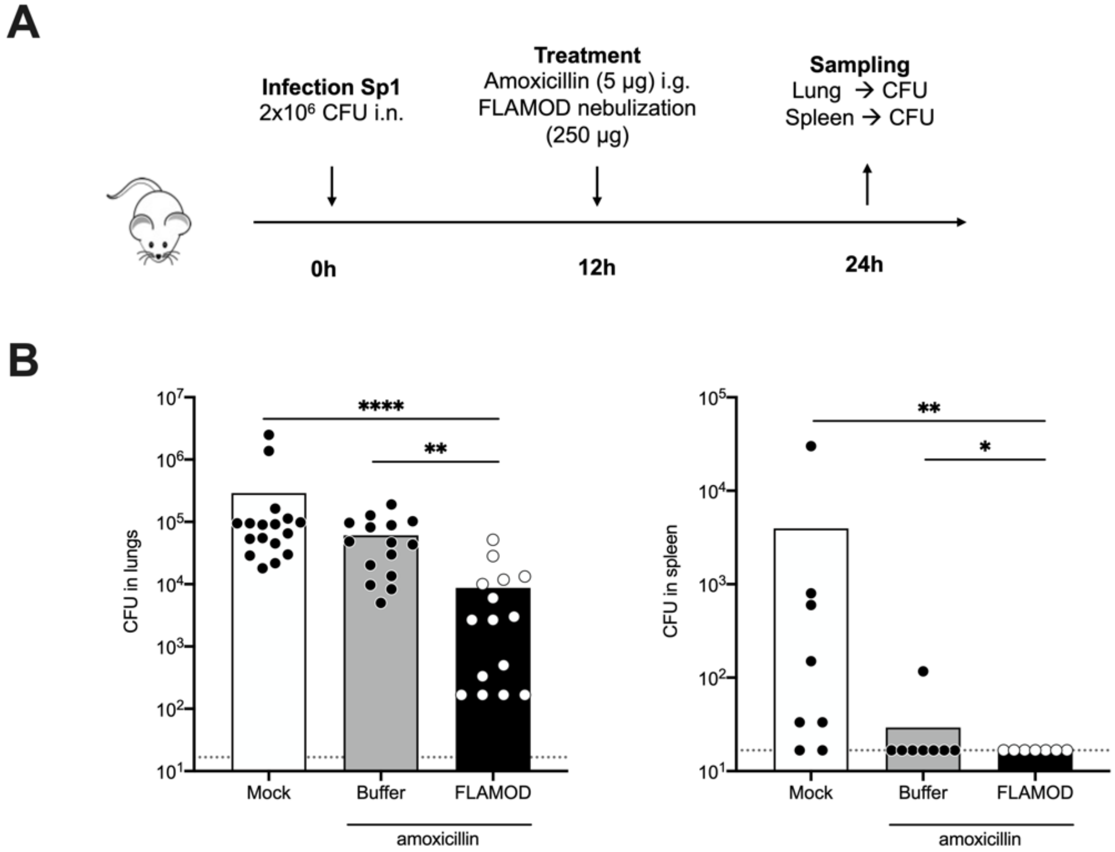


**Supplementary Figure 5. FLAMOD inhalation efficacy in a mouse model of primary pneumococcal pneumonia.** BALB/c mice were infected intranasally with $2x10^6$ CFU of *S. pneumoniae* serotype 1. Animals were treated 12 hours post infection by nebulization with 1 ml of FLAMOD at 250 µg/ml in NaPi buffer combined with intragastric administration of 5 µg amoxicillin. Mock- and amoxicillin alone-treated animals received NaPi buffer and water, and NaPi buffer and amoxicillin, respectively. Lungs (n=15-17 per condition) and spleen (n=7-8 per condition) were sampled 12 h post-administration for CFU. **(A)** Experimental design. **(B)** CFU in the lungs and spleen of infected and treated mice. Dashed lines indicate limit of quantification. Statistical differences between the groups were analyzed by One-way ANOVA (*p<0.05, **p<0.01, ****p<0.0001).
