## Supplementary Table 1 for "Targeted delivery of flagellin by nebulization offers optimized respiratory immunity and defense against pneumococcal pneumonia"

**Supplementary Table 1**. **Forward and reverse primer sequences for qPCR**

| **Target gene** | **Forward primer** | **Reverse primer** |
| --- | --- | --- |
| *Actb* | CGTCATCCATGGCGAACTG | GCTTCTTTGCAGCTCCTTCGT |
| *B2m* | TGGTCTTTCTGGTGCTTGTC | GGGTGGCGTGAGTATACTTGAA |
| *Ccl20* | TTTTGGGATGGAATTGGACAC | TGCAGGTGAAGCCTTCAACC |
| *Cxcl2* | CCCTCAACGGAAGAACCAAA | CACATCAGGTACGATCCAGGC |
| *Il6* | TCTAATTCATATCTTCAACCAAGAGG | TGGTCCTTAGCCACTCCTTC |
